# Population Genetic Structure of *Ariolimax columbianus* around Corvallis, Oregon, USA

**DOI:** 10.64898/2026.07.21.739958

**Authors:** Maximillian Brune, Dana K. Howe, Rory J. Mc Donnell, Dee Denver

## Abstract

Urbanization has been previously shown to affect evolution of plants and animals by restricting gene flow, increasing genetic drift, and causing divergent selection between rural and urban populations, however, the effect of urbanization on terrestrial gastropod evolution has only been minimally researched. Most research on terrestrial gastropods primarily focusing on invasive pest species, and little attention is given to native species like *Ariolimax columbianus* (Pacific banana slug). There is no current published research on the phylogeography of *A. columbianus.* In this study, surface epithelial cells from 66 *Ariolimax columbianus* individuals were collected from 11 locations around Corvallis, Oregon, and one site in northern California. These samples were amplified and sequenced at two loci: cytochrome oxidase 1 (CO1) and the internal transcribed spacer 2 (ITS-2). This data was used to generate maximum-likelihood phylogenetic trees to better understand the population genetics of these local native gastropods. The CO1 phylogenetic analysis showed two major haplotypes around the city of Corvallis – northwestern and southern, while the ITS-2 phylogenetic analysis showed three general haplotypes around the city – northern, western, and southern. The results in this study show the presence of genetic breaks between many of these populations, possibly due to a combination of factors including anthropogenic fragmentation, natural barriers, and the low active dispersal ability of *A. columbianus*. However, the extent that each of those factors has on disrupting gene flow and explaining patterns of genetic variation in this species requires further research. The present study acts as a foundation for the development of future research on banana slug phylogenetics and population genetic structure.

## Introduction

As the world’s population grows, urbanization continues to predominate with over half of the world’s population currently residing in urban environments (Miles et al., 2019). This urbanization has previously been shown to affect natural selection, promote a higher mutation rate due to increased pollution, and alter natural gene flow between different populations (Rivkin et al., 2019). Urban developments and landscape alterations can cause both habitat loss and habitat fragmentation. The loss of a habitat often results in decreased fitness of a species as it is driven to habitats that are suboptimal for its growth and reproduction (Nicolai & Ansart, 2017). Meanwhile, fragmentation may lead to increased isolation of specific populations and therefore reduced gene flow if organisms are unable to migrate between these populations. Genetic drift would subsequently occur and lead to higher levels of differentiation between these populations, particularly in the smaller populations that arise after fragmentation. Further, if the populations are isolated in differing environments even across the same city, selective pressures may further contribute to increasing genetic variation.

This effect would likely be especially prevalent for gastropods, the largest group within the Mollusca phylum, making up an estimated 80% of all extant molluscan taxa (Bieler, 1992; Razkin et al., 2015). Within Mollusca, the clade Stylommatophora (Gastropoda: Pulmonata) composes the majority of the extant terrestrial mollusc species living in a wide variety of distinct terrestrial habitats and microhabitats, including grasslands, river embankments, decaying wood, and leaf litter in forests (Leonard & Cordoba-Aguilar, 2010). The genus *Ariolimax* (banana slugs) within Stylommatophora includes several large species of slugs endemic to the west coast of North America (Montroni et al., 2019). Among these species, the Pacific banana slug, *Ariolimax columbianus*, has a large range spanning from the coast of central California to southeastern Alaska, and is one of the largest terrestrial slugs in North America, reaching lengths of up to 260 mm when fully extended (von Proschwitz et al., 2017). There are several different color variations that have complicated prior taxonomic classifications, as *A. columbianus* individuals within the same populations have been found to be white, yellow, brown, or black, and they can be either monochromatic or maculated (Pearson et al., 2006). While there are several other species within *Ariolimax*, *A. columbianus* is the only one with a range known to extend outside California.

Current literature on gastropods has primarily focused on invasive pest slugs rather than native species such as *A. columbianus*. This is partly due to the prevalence of invasive slugs, as slugs represent one-third of the entire introduced fauna in North America while only representing approximately 7% of the native fauna (Nekola, 2014). Non-native pest slugs cause millions of dollars in agricultural damage every year in Oregon, increasing researchers and farmers attempts to find methods for controlling these pest slug populations (Mc Donnell et al., 2020). Some of these methods involve the use of slug-parasitic nematodes and molluscicides to kill these pests, however, native slug conservation is a particularly important issue, especially regarding the controversial history of numerous failed molluscan management strategies (Christensen et al., 2021). Therefore, further research into native slug species such as *A. columbianus* is warranted to expand the currently limited knowledge on these slugs.

Previous research on *A. columbianus* has largely focused on examining feeding and reproductive behaviors. They are known to feed on a variety of plant species and organic detritus, and are proposed to have a role in seed dispersal (Gervais et al., 1998; Rodriguez-Cabal et al., 2015). Much of the previous research that has been done on banana slugs has involved the taxonomic classifications of *Ariolimax* using reproductive system anatomy (Mead, 1943), as well as the use of *Ariolimax* to study sexual conflict and mating systems (Leonard, 1991) because of their role as simultaneous hermaphrodites and the intriguing behavior of gnawing off the penis after copulation. Although dispersal has been studied in other gastropod species (Aubry et al., 2006), little is known about active and passive dispersal patterns in *A. columbianus*, and there is no current published research on its phylogeography. Other large-scale phylogeographic studies involving other species have been done in the Pacific Northwest region of North America, which showed evidence of genetic breaks between north and south populations of several species of plants which was suspected to have been caused by past glaciation affecting the genetic structure of the Pacific Northwest flora and fauna (Soltis et al., 1997). A similar study was done on the arionid slug *Prophysaon coeruleum* showing genetic variation likely caused by past geological events (Wilke & Duncan, 2004). However, the local evolutionary responses of terrestrial gastropods such as *A. columbianus* to urbanization and habitat fragmentation has not been studied in depth (Johnson & Munshi-South, 2017; Vendetti et al., 2018). While these large-scale studies have been explored in-depth, the present study takes a smaller scaled approach by seeking to assess the local genetic population structure of *A. columbianus* around the city of Corvallis, Oregon.

This study aims to further the understanding of the population genetics of these native gastropods by comparing sequence information from a portion of the mitochondrial cytochrome oxidase 1 (CO1) and the nuclear internal transcribed spacer 2 (ITS-2) DNA from local *A. columbianus* around Corvallis. The central hypothesis tested is that the city of Corvallis, Oregon disrupts gene flow between isolated populations of *A. columbianus*. Given previous research on the effects of urbanization on habitat fragmentation, genetic drift, and natural selection, it was predicted that DNA sequence analysis would reveal that populations separated by the city would demonstrate distinct haplotypes. Genetic variation between separated and isolated populations of *A. columbianus* would suggest that the city of Corvallis indeed acts as a barrier to gene flow by preventing the dispersal of local banana slugs between populations.

## Methods and Materials

### Sampling sites and DNA collection

To test the above-stated hypothesis, a set of locations for sampling was first determined (Figure 1). During the first round of sampling that took place from February 2021 to November 2021, seven locations were selected within a ten-mile radius of Oregon State University in Corvallis, Oregon. During the second round of sampling from September 2022 to November 2022, another five locations were sampled including a distant population from Humbolt County, California. In total, 66 samples were collected from 12 unique sites.

**Figure 1:**
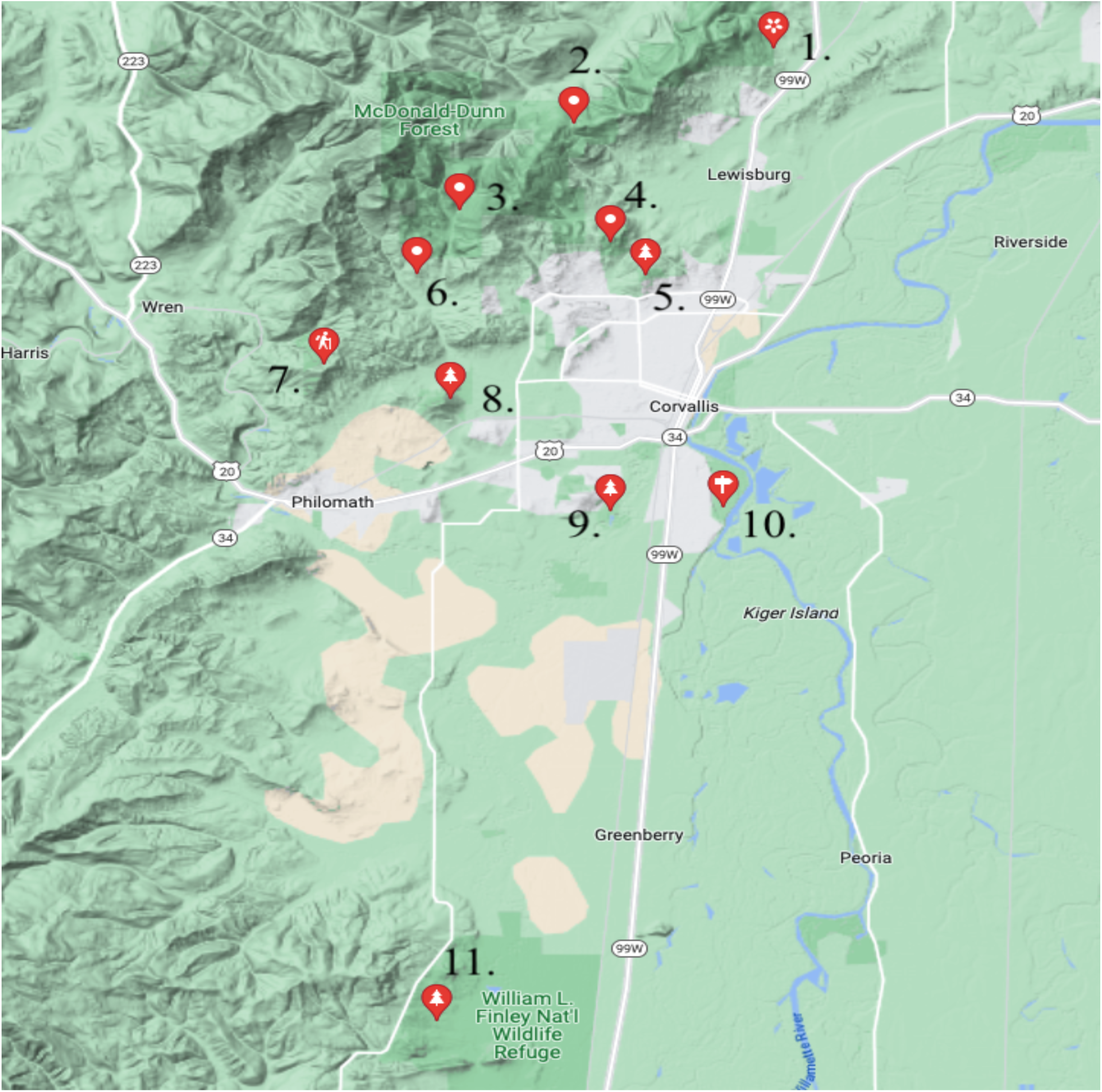
Map of the 11 sampling locations around Corvallis, Oregon. Listed from north to south, they include: Peavy Arboretum (PA), Lewisburg Saddle (LS), Oak Creek (OC), Chip Ross Park (CR), Brandis City Natural Area (BC), Private Residence (PR),Fitton Green Natural Area (FG), Bald Hill Natural Area, Mary’s River Natural Area, Willamette Park, and William Finley National Wildlife Refuge (WF). An additional sampling location in Humboldt County (HC) in northern California is not shown in this figure.

Most of the literature that exists on terrestrial slugs is focused on agricultural pest species, so many past methods of DNA collection do not place an emphasis on the conservation of these populations and instead are intrusive and harmful to the slug (Morinha et al., 2014). However, given the present study’s focus on benign native banana slugs, it was important to minimize any harmful effects from the sampling process. Epithelial cells on the slugs’ bodies were collected by scraping a sterile cotton-tipped swab ten times along the mantle, all the way down the body of the slug. These swabs were then placed into 1.5 mL Eppendorf tubes and placed on ice until sampling at the location was finished. Swabs were then stored at -20°C until DNA extraction. At the time of collection, each slug with some exceptions, was indexed with a picture and coordinates on the citizen science application iNaturalist (Supplemental Table 1).

### Two protocols for DNA extraction

Two different protocols were carried out for DNA extraction. The first protocol, used on samples collected in 2021, was a slight modification of the lysis buffer method from Williams et al. (1992). This method proved to have a low success rate of the subsequent PCR and sequencing (only 38.2% of samples successfully amplified and sequenced), so the extraction protocol was modified and adapted to follow the conventional salting out protocol from Morinha et al. (2014). This new modified protocol was used for the rest of the samples collected in 2022.

### PCR and Sanger sequencing

For PCRs amplifying the CO1 gene, primers LCO1490 and HCO2198 were used (Folmer et al., 1994). The PCRs amplifying the ribosomal gene cluster (including the partial 5.8S gene, the partial 28S gene, and the entire ITS-2 region between them), primers LSU-1 and LSU-3 were used (Wade et al., 2006). All reactions were run on a 1.5% agarose gel to assess for successful amplification and purified using a nano beading procedure (Elkin et al., 2001). Direct-end Sanger sequencing was done using BigDye Terminator v3.1 Cycle Sequencing Kit (Applied Biosystems) on an ABI 3730 capillary machine at the Oregon State University Center for Quantitative Life Sciences. DNA sequences generated for this study were submitted to Genbank under accession numbers (to be deposited).

### Phylogenetic analysis

Sequence results were quality checked and aligned using MEGA X (Kumar et al., 2018). Before phylogenetic analysis was performed, the model selection tool in MEGA X was employed to find the most appropriate model with the lowest BIC for each alignment. For the 529 aligned base pairs of the CO1 locus, the General Time Reversible model with Gamma distribution (Nei & Kumar, 2000) was used to generate a maximum-likelihood phylogeny with 1,000 bootstrap replications. In addition to the 45 CO1 sequences from newly sampled slugs, five *Ariolimax columbianus* CO1 sequences found in GenBank were added to the analysis (see Figure 2 for accession numbers). Although three of the sequences were labeled as *Stylommatophora sp.* and one was labeled as *Arionidae sp.* (all from Vancouver, BC), photos of those individuals were consistent with the morphology of *A. columbianus* and they shared a high percent identity on BLAST search. Given the known range of the species within the *Ariolimax* genus can extend outside of California, it is likely that these individuals represent *A. columbianus* and were thus included in the phylogenetic analysis. The five outgroups were also added for the CO1 phylogeny: *Mysticarion obscurior* (Hyman et al., 2017), *Asperitas cidaris, Asperitas stuartiae, Asperitas inquinata,* and *Asperitas bimaensis* (Köhler et al., 2020).

**Figure 2:**
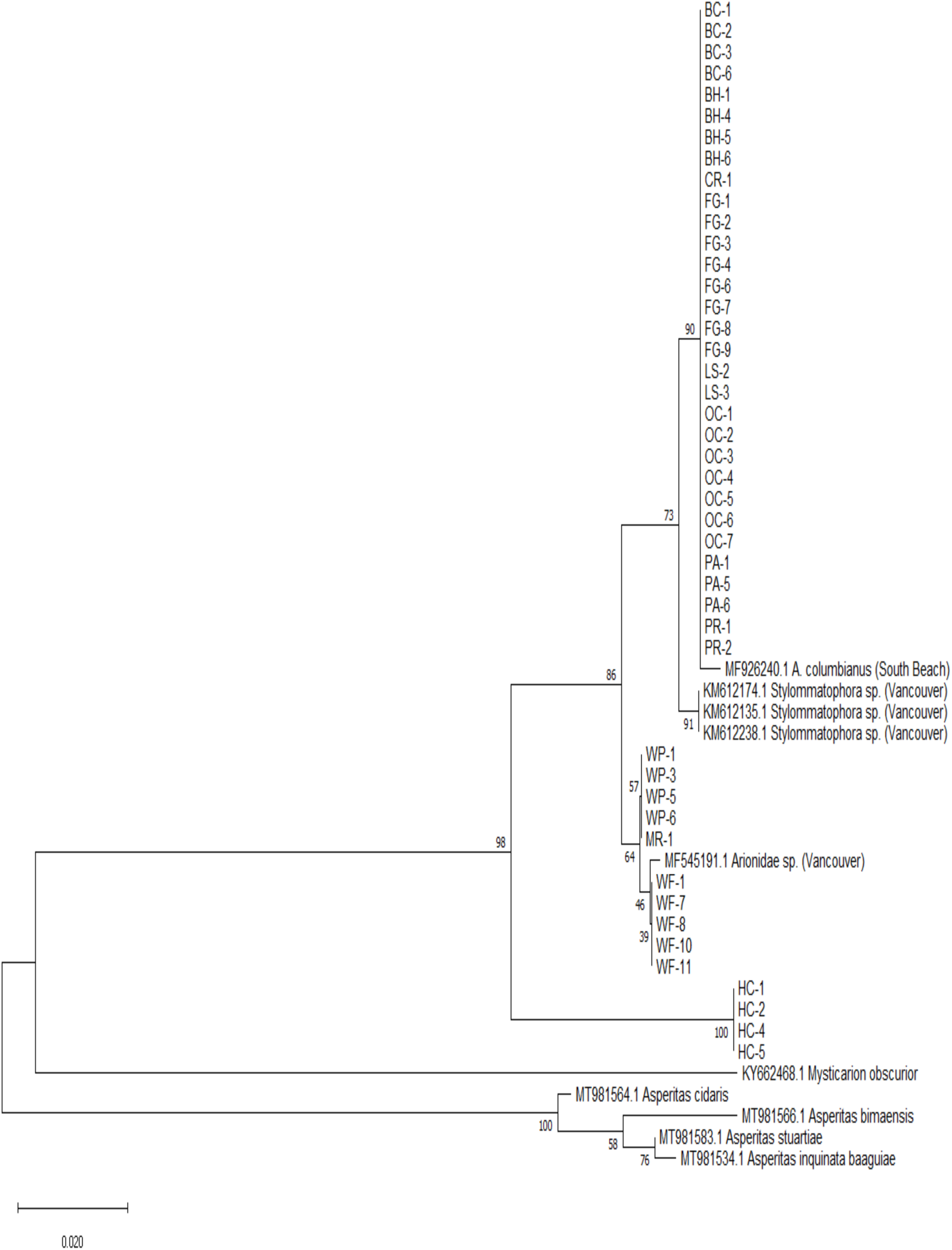
Maximum likelihood phylogeny of 529 bp of the mitochondrial CO1 gene. A total of 55 CO1 sequences were included in the phylogeny, 45 new sequences from slugs sampled for this study, five *Ariolimax columbianus* sequences obtained from GenBank (although they had not been initially identified as *A. columbianus*) and five outgroups. Node values represent the maximum likelihood value and the scale bar represents 0.020 nucleotide substitutions per site.

For the analysis of the 820 aligned base pairs from the ITS-2 locus, the Tamura-Nei model with Gamma distribution (Tamura & Nei, 1993) was used to create the maximum-likelihood phylogeny with 1,000 bootstrap replications. A total of of 53 ITS-2 sequences were included, 48 new sequences from slugs from this study and five outgroups available in Genbank: *Limax conemenosi* (Giusti et al., 2021), *Candaharia rutellum*, *Trigonochlamys imitatrix, Aegopinella nitidula* (Neiber et al., 2020) and *Oligolimax annularis* analysis (see Figure 3 for accession numbers).

**Figure 3:**
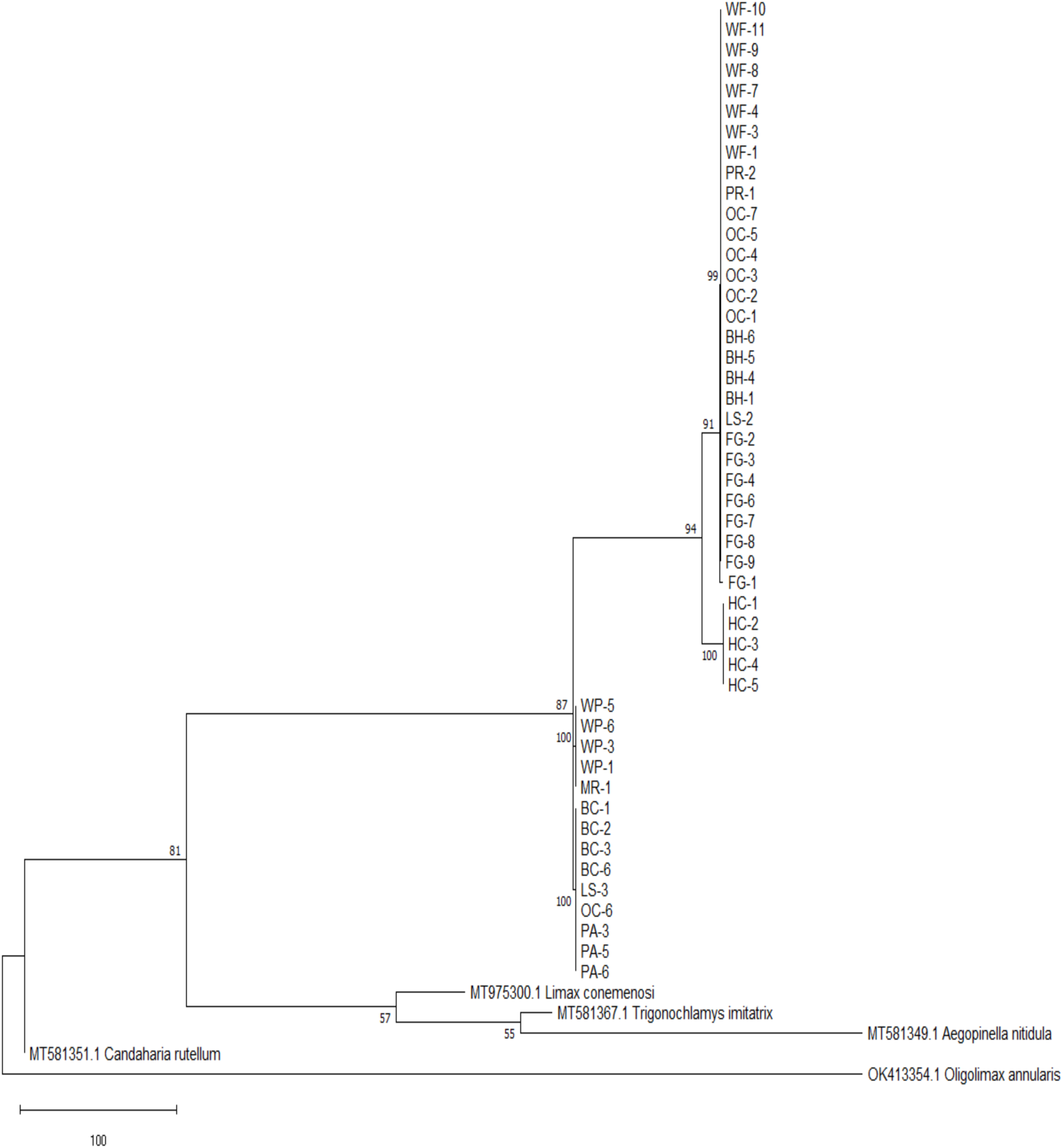
Maximum likelihood phylogeny of 820 bp of the internal transcribed spacer (ITS-2) with flanking regions of 5.8S and 28S. There were a total of 53 ITS-2 sequences included in the phylogeny, 48 sequences from slugs that had been directly sequenced for the present study and with five outgroups. Node values represent the maximum likelihood value and the scale bar represents 100 nucleotide substitutions per site.

## Results

Out of the 66 epithelial cell samples collected from *A. columbianus* from 12 different locations, 45 were successfully sequenced at the CO1 locus and 48 were sequenced at the ITS-2 locus, with 43 that were sequenced at both loci. This discrepancy was largely due to the inefficient initial DNA extraction method, and its modification made later samples more successful.

The phylogeny for the mitochondrial CO1 locus (Figure 2) shows 31 samples from the eight locations to the northwest of Corvallis, Oregon form a monophyletic group with strong support, while the 10 samples from the locations south of the city [Mary’s River Natural Area (MR), Willamette Park (WP), and William L. Finley National Wildlife Refuge (WF)] form a separate group. The Humboldt County location from northern California also forms a unique monophyletic group with strong support, and those slugs were shown to be the most distantly related among any of the sampled slugs.

The phylogeny for the nuclear ITS-2 locus (Figure 3) overall shows strong support for four major clades that can largely be described geographically. Twenty-eight slug sequences from western Corvallis collected from a private residence (PR), Oak Creek (OC), Bald Hill (BH), Fitton Green (FG), and WF locations form a distinct clade, with the exception of one slug from FG. Further, the northern Corvallis populations from Peavy Arboretum (PA) and Brandis City Natural Area (BC) form a clade, the southern Corvallis slug samples from MR and WP form a clade, and the Humboldt County (HC) slugs form their own clade (Figure 3). Of note, two sampling sites – Lewisburg Saddle (LS) and OC – are shown to be polyphyletic, with individuals in both the northern and western groups. In the case of LS, there were two samples sequenced with one grouping in the western clade and the other grouping in the northern clade. For OC, only one of the seven sequences grouped with the northern clade, while the other six were with the western clade. These two sites are in close proximity to each other and are located within the same forest to the northwest of the city, so the overlap is not entirely surprising. Additionally, the polytomies observed in this phylogeny suggesting there is not enough genetic data to elucidate more precise relationships among the northern, western (plus WF), and southern Corvallis groups. However, the western group was shown to be more closely related to the HC slugs than to any of the other clades.

## Discussion

The data presented in this study provide evidence of genetic variation among geographically isolated populations. The CO1 phylogeny shows the samples collected in the study separating into three major clades that are able to be described geographically (Figure 1) – northwestern Corvallis, southern Corvallis including WF, and the HC group in northern California. Meanwhile, the ITS-2 phylogeny shows four different clades, with the presence of unique northern and western clades compared to the single northwestern group of the CO1 phylogeny. Overall, the ITS-2 analysis shows a greater amount of genetic variation between sites, but the overall evolutionary story is somewhat similar with divergence between populations that are separated by the city of Corvallis (with one exception of the WF slugs in the ITS-2 phylogeny).

This evidence suggests that there is a disruption in gene flow between these different populations, allowing for genetic drift and possibly natural selection to cause molecular divergence. However, the extent to which anthropogenic fragmentation, natural factors, or simply the limited active dispersal abilities of *A. columbianus* affect this disruption in gene flow is still uncertain. The MR and WP locations just south of the city, although are only about 1.5 miles away from each other, are separated by a river, a busy highway, and many industrial and residential buildings. Regardless, they shared the same haplotype in both phylogenies while populations separated by just a couple miles of mostly old growth forest separate into different haplotypes (northern versus western clades) in the ITS-2 phylogeny. Further studies with greater sampling density and objective measures of population isolation and factors involved in disrupting gene flow would be required to elucidate the effect and strength of those factors.

Additionally, it has been shown that past geological and glaciation events have influenced the distribution of genetic variation in various plants and animals in the Pacific Northwest (Soltis et al., 1997; Wilke & Duncan, 2004). Further work would need to be done using molecular clock rates to date the divergence and identify if any historical processes contributed to the genetic variation demonstrated in this study. Finally, calculations regarding effective population size and exploration of the effect of possible natural selection on these populations would be able to provide further knowledge about the evolutionary forces involved in creating the divergence seen between the sampled populations.

The phylogeographic pattern demonstrated at the ITS-2 locus is somewhat surprising, particularly because the slugs from the locations in western Corvallis – PR, BH, OC, and FG – shared the same clade as slugs from the WF site and were more divergent with populations from other sites that are in closer proximity. Given the distance, presence of anthropogenic structures and farmland, and the low active dispersal rate of terrestrial gastropods such as *A. columbianus* (Aubry et al., 2006), it was unexpected for these sampling sites to share a most recent common ancestor while closer sites are more divergent. Similarly, the slugs from the western area were shown to be more closely related to the HC slugs from northern California in the ITS-2 phylogenetic tree than the northern or southern Corvallis clades, despite being located over 300 miles away. The reason for these findings is unclear, although one possibility is that slugs native to one site were passively transported to the other location via facilitation by humans, animals, or other natural factors. Several *A. columbianus* individuals have been recorded in Europe despite the lack of any known established populations of *A. columbianus* there (von Proschwitz et al., 2017), with evidence that they were accidentally imported to Swedish flower shops along with ornamental plants from the Pacific Northwest. Given these records of intercontinental transport of *A. columbianus*, as well as previous evidence of the passive dispersal of other terrestrial slugs and snails (Aubry et al., 2006; Gittenberger, 2012; Ożgo et al., 2016; Simonová et al., 2016), it is not unlikely that passive dispersal of slugs from one location to the other could be responsible for this finding, especially since *A. columbianus* are capable of self-fertilization, and therefore a new population could theoretically be started from the transport of just one slug.

Another future direction for research involves the greater incorporation of citizen science initiatives such as iNaturalist. The smartphone application and website is an increasingly popular citizen science tool that has shown great promise and utility, particularly for projects involving ecology, conservation, and biodiversity (Mesaglio & Callaghan, 2021). It has been used in Australia to record the diversity and distribution of sea slugs (Smith & Nimbs, 2022); users on the app have also observed a rare, threatened species of Philippine Bumble Bee, *Bombus irisanensis*, that had previously not been reported since the 1990s (Wilson et al., 2020); and users documented the first occurrence records of five introduced terrestrial gastropod species in southern California (Vendetti et al., 2018). The use of citizen science initiatives allows for extraordinary levels of data collection far beyond what a single research team could collect themselves, providing the opportunity for large-scale studies regarding ecology, biodiversity, and the distribution of not only *A. columbianus* but also invasive pest slugs that cause millions of dollars in agricultural damage every year (Mc Donnell et al., 2020).

## Supplemental Material

**Supplemental Figure 1:**
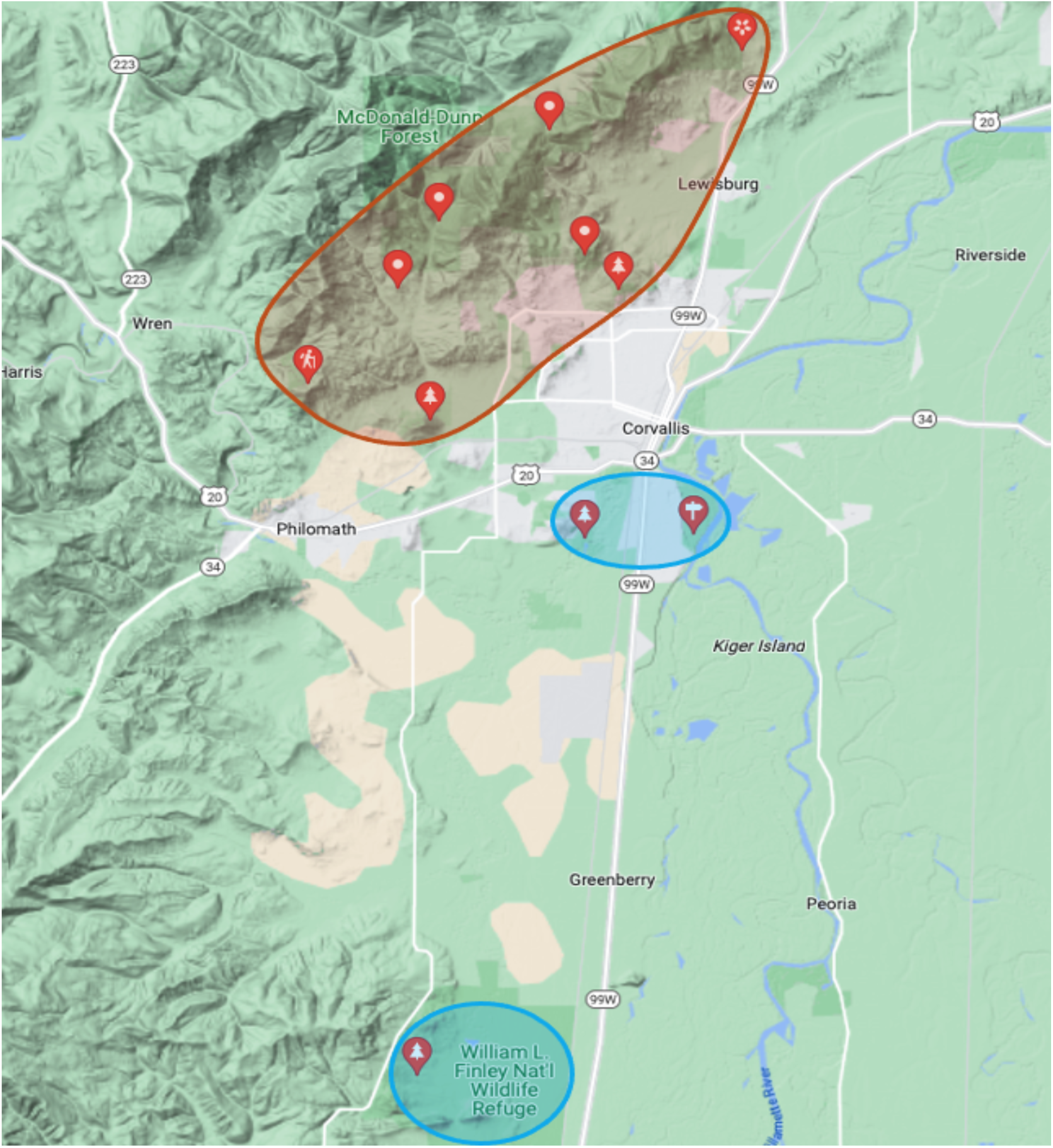
Map of Corvallis, Oregon showing the sampling locations (marked with red pins) and the geographic distribution of the clades demonstrated by the CO1 phylogeny. Orange represents the northwestern clade and light blue represents the southern clade consisting of the William L. Finley, Mary’s River, and Willamette Park locations. The clade consisting of the slugs from Humboldt County in northern California are not pictured.

**Supplemental Figure 2:**
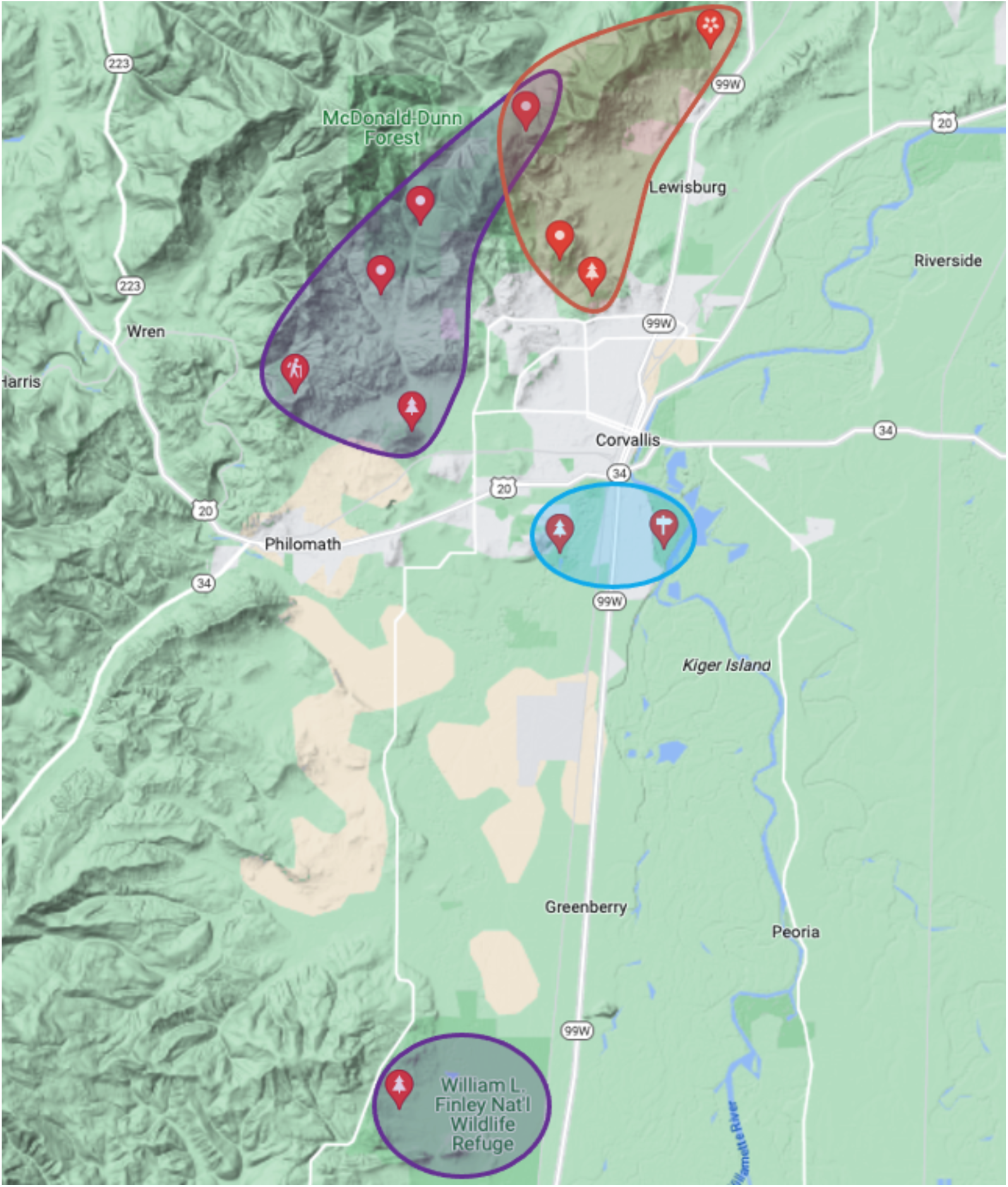
Map of Corvallis, Oregon showing the sampling locations (marked with red pins) and the geographic distribution of the clades demonstrated by the ITS-2 phylogeny. Orange represents the northern clade, light blue represents the southern Corvallis clade, and purple represents the western clade (plus slugs from the William L. Finley National Wildlife Refuge). The clade consisting of the slugs from Humboldt County in northern California are not pictured.

**Supplementary Table 1.** *A. columbianus* sample information. List of the banana slug samples along with their collection sites, dates and iNaturalist links.

| Location | Slug Identifier | Collection Date | iNaturalist Link |
| --- | --- | --- | --- |
| Private Residence | PR-1 | 2/23/21 | https://www.inaturalist.org/observations/116566279 |
|  | PR-2 |  | https://www.inaturalist.org/observations/116565959 |
|  | PR-3 |  | N/A |
| Peavy Arboretum | PA-1 | 4/18/21 | https://www.inaturalist.org/observations/74402047 |
|  | PA-2 | 11/18/21 | https://www.inaturalist.org/observations/101412283 |
|  | PA-3 |  | https://www.inaturalist.org/observations/101417074 |
|  | PA-4 |  | https://www.inaturalist.org/observations/101417080 |
|  | PA-5 |  | https://www.inaturalist.org/observations/101417085 |
|  | PA-6 |  | N/A |
| Brandis City Natural Area | BC-1 | 4/23/21 | https://www.inaturalist.org/observations/74987762 |
|  | BC-2 |  | https://www.inaturalist.org/observations/74988572 |
|  | BC-3 |  | https://www.inaturalist.org/observations/74989397 |
|  | BC-4 |  | https://www.inaturalist.org/observations/74991289 |
|  | BC-5 |  | N/A |
|  | BC-6 |  | https://www.inaturalist.org/observations/74991086 |
|  | BC-7 |  | https://www.inaturalist.org/observations/74991015 |
| Willamette Park | WP-1 | 4/30/21 | https://www.inaturalist.org/observations/76005729 |
|  | WP-2 | 5/20/21 | https://www.inaturalist.org/observations/79544644 |
|  | WP-3 |  | https://www.inaturalist.org/observations/79546198 |
|  | WP-4 |  | https://www.inaturalist.org/observations/79547423 |
|  | WP-5 | 11/12/22 | https://www.inaturalist.org/observations/141835645 |
|  | WP-6 |  | https://www.inaturalist.org/observations/141839403 |
| Bald Hill Natural Area | BH-1 | 5/6/21 | https://www.inaturalist.org/observations/77691514 |
|  | BH-2 |  | https://www.inaturalist.org/observations/77692388 |
|  | BH-3 |  | https://www.inaturalist.org/observations/77693420 |
|  | BH-4 | 10/27/22 | https://www.inaturalist.org/observations/140174597 |
|  | BH-5 |  | https://www.inaturalist.org/observations/140182872 |
|  | BH-6 |  | https://www.inaturalist.org/observations/140217050 |
| Mary's River Natural Area | MR-1 | 5/27/21 | https://www.inaturalist.org/observations/116567032 |
| Lewisburg Saddle | LS-1 | 11/12/21 | https://www.inaturalist.org/observations/100957116 |
|  | LS-2 | 10/9/22 | https://www.inaturalist.org/observations/138146344 |
|  | LS-3 |  | https://www.inaturalist.org/observations/138147339 |
| William L. Finley National Wildlife Reserve | WF-1 | 9/28/22 | https://www.inaturalist.org/observations/136847877 |
|  | WF-2 |  | https://www.inaturalist.org/observations/136848498 |
|  | WF-3 |  | https://www.inaturalist.org/observations/136853564 |
|  | WF-4 |  | https://www.inaturalist.org/observations/136853652 |
|  | WF-5 | 9/29/22 | https://www.inaturalist.org/observations/136977367 |
|  | WF-6 |  | https://www.inaturalist.org/observations/136973548 |
|  | WF-7 |  | https://www.inaturalist.org/observations/136974672 |
|  | WF-8 |  | https://www.inaturalist.org/observations/136976526 |
|  | WF-9 |  | https://www.inaturalist.org/observations/136977378 |
|  | WF-10 | 9/30/22 | https://www.inaturalist.org/observations/137094535 |
|  | WF-11 |  | https://www.inaturalist.org/observations/137094554 |
| Fitton Green Natural Area | FG-1 | 10/10/22 | https://www.inaturalist.org/observations/138270339 |
|  | FG-2 |  | https://www.inaturalist.org/observations/138271262 |
|  | FG-3 |  | https://www.inaturalist.org/observations/138271313 |
|  | FG-4 |  | https://www.inaturalist.org/observations/138273802 |
|  | FG-5 |  | https://www.inaturalist.org/observations/138275471 |
|  | FG-6 |  | https://www.inaturalist.org/observations/138280887 |
|  | FG-7 |  | https://www.inaturalist.org/observations/138280894 |
|  | FG-8 |  | https://www.inaturalist.org/observations/138281040 |
|  | FG-9 |  | https://www.inaturalist.org/observations/138281066 |
| Oak Creek | OC-1 | 10/18/22 | https://www.inaturalist.org/observations/139655390 |
|  | OC-2 |  | https://www.inaturalist.org/observations/139655400 |
|  | OC-3 |  | https://www.inaturalist.org/observations/139655413 |
|  | OC-4 |  | https://www.inaturalist.org/observations/139655429 |
|  | OC-5 |  | https://www.inaturalist.org/observations/139655441 |
|  | OC-6 |  | https://www.inaturalist.org/observations/139655463 |
|  | OC-7 |  | https://www.inaturalist.org/observations/139655473 |
| Chip Ross Park | CR-1 | 10/22/22 | https://www.inaturalist.org/observations/139660436 |
| Humboldt County, California | HC-1 | 7/14/22 | N/A |
|  | HC-2 |  |  |
|  | HC-3 |  |  |
|  | HC-4 |  |  |
|  | HC-5 |  |  |
|  | HC-6 |  |  |

## Acknowledgements

We thank Casey Richart, Emily Taylor, and Indira Kulkarni for feedback and comments on the research and manuscript. Research funding was provided by Oregon State University.

## References

Aubry, S., Labaune, C., Magnin, F., Roche, P., & Kiss, L. (2006). Active and Passive Dispersal of an Invading Land Snail in Mediterranean France. Journal of Animal Ecology, 75(3), 802–813.

Bieler, R. (1992). Gastropod Phylogeny and Systematics. Annual Review of Ecology and Systematics, 23(1), 311–338. 10.1146/annurev.es.23.110192.001523

Elkin, C. J., Richardson, P. M., Fourcade, H. M., Hammon, N. M., Pollard, M. J., Predki, P. F., Glavina, T., & Hawkins, T. L. (2001). High-Throughput Plasmid Purification for Capillary Sequencing. Genome Research, 11(7), 1269–1274. 10.1101/gr.167801

Folmer, O., Black, M., Hoeh, W., Lutz, R., & Vrijenhoek, R. (1994). DNA primers for amplification of mitochondrial cytochrome c oxidase subunit I from diverse metazoan invertebrates. Molecular Marine Biology and Biotechnology, 3(5), 294–299.

Gervais, J. A., Traveset, A., & Willson, M. F. (1998). The Potential for Seed Dispersal by the Banana Slug (Ariolimax columbianus). The American Midland Naturalist, 140(1), 103–110. 10.1674/0003-0031(1998)140[0103:TPFSDB]2.0.CO;2

Gittenberger, E. (2012). Long-distance dispersal of molluscs: ‘Their distribution at first perplexed me much.’ Journal of Biogeography, 39(1), 10–11. 10.1111/j.1365-2699.2011.02638.x

Giusti, F., Lesicki, A., Benocci, A., Pieńkowska, J. R., & Manganelli, G. (2021). Weltersia obscura, a new slug from the island of Montecristo (Tuscan Archipelago, Italy): A hitherto undiscovered endemic or a recent alien? (Mollusca, Pulmonata, Limacidae). Systematics and Biodiversity, 19(7), 648–664. 10.1080/14772000.2021.1908442

Hyman, I., Lamborena, I., & Köhler, F. (2017). Molecular phylogenetics and systematic revision of the south-eastern Australian Helicarionidae (Gastropoda, Stylommatophora). Contributions to Zoology Bijdragen Tot de Dierkunde, 86, 51–95. 10.1163/18759866-08601004

Johnson, M. T. J., & Munshi-South, J. (2017). Evolution of life in urban environments. Science, 358(6363), eaam8327. 10.1126/science.aam8327

Köhler, F., Criscione, F., Hallan, A., Hyman, I., & Kessner, V. (2020). Lessons from Timor: Shells are poor taxonomic indicators in Asperitas land snails (Stylommatophora, Dyakiidae). Zoologica Scripta, 49(6), 732–745. 10.1111/zsc.12449

Kumar, S., Stecher, G., Li, M., Knyaz, C., & Tamura, K. (2018). MEGA X: Molecular Evolutionary Genetics Analysis across Computing Platforms. Molecular Biology and Evolution, 35(6), 1547–1549. 10.1093/molbev/msy096

Leonard, J. (1991). Sexual conflict and the mating systems of simultaneously hermaphrodite gastropods. American Malacological Bulletin, 9, 45–58.

Leonard, J. L., Córdoba-Aguilar A., & Baur, B. (2010). Stylommatophoran Gastropods. In The evolution of primary sexual characters in animals (pp. 197–217). essay, Oxford University Press.

Mc Donnell, R. J., Colton, A. J., Howe, D. K., & Denver, D. R. (2020). Lethality of four species of Phasmarhabditis (Nematoda: Rhabditidae) to the invasive slug, Deroceras reticulatum (Gastropoda: Agriolimacidae) in laboratory infectivity trials. Biological Control, 150, 104349. 10.1016/j.biocontrol.2020.104349

Mead, A. R. (1943). Revision of the Giant West Coast Land Slugs of the Genus Ariolimax Moerch (Pulmonata: Arionidae). The American Midland Naturalist, 30(3), 675–717. 10.2307/2421208

Mesaglio, T., & Callaghan, C. T. (2021). An overview of the history, current contributions and future outlook of iNaturalist in Australia. Wildlife Research, 48(4), 289–303. 10.1071/WR20154

Miles, L. S., Rivkin, L. R., Johnson, M. T. J., Munshi-South, J., & Verrelli, B. C. (2019). Gene flow and genetic drift in urban environments. Molecular Ecology, 28(18), 4138– 4151. 10.1111/mec.15221

Montroni, D., Zhang, X., Leonard, J., Kaya, M., Amemiya, C., Falini, G., & Rolandi, M. (2019). Structural characterization of the buccal mass of Ariolimax californicus (Gastropoda; Stylommatophora). PLOS ONE, 14(8), e0212249. 10.1371/journal.pone.0212249

Morinha, F., Travassos, P., Carvalho, D., Magalhães, P., Cabral, J. A., & Bastos, E. (2014). DNA sampling from body swabs of terrestrial slugs (Gastropoda: Pulmonata): a simple and non-invasive method for molecular genetics approaches. Journal of Molluscan Studies, 80(1), 99–101. 10.1093/mollus/eyt045

Nei M., & Kumar S. (2000). Molecular Evolution and Phylogenetics. Oxford University Press, *New York*.

Neiber, M. T., Walther, F., & Hausdorf, B. (2020). Phylogenetic relationships of ghost slugs (Selenochlamys) and overlooked instances of limacization in Western Palaearctic Limacoidei (Gastropoda: Stylommatophora). Molecular Phylogenetics and Evolution, 151, 106897. 10.1016/j.ympev.2020.106897

Nekola, J. C. (2014). Overview of the North American Terrestrial Gastropod Fauna. American Malacological Bulletin, 32(2), 225. 10.4003/006.032.0203

Nicolai, A., & Ansart, A. (2017). Conservation at a slow pace: Terrestrial gastropods facing fast-changing climate. Conservation Physiology, 5(1), cox007. 10.1093/conphys/cox007

Ożgo, M., Örstan, A., Kirschenstein, M., & Cameron, R. (2016). Dispersal of land snails by sea storms. Journal of Molluscan Studies, 82(2), 341–343. 10.1093/mollus/eyv060

Pearson, A. K., Pearson, O. P., & Ralph, P. L. (2005). Growth and Activity Patterns in a Backyard Population of the Banana Slug. The Veliger, 48(3), 143–150.

Razkin, O., Gómez-Moliner, B. J., Prieto, C. E., Martínez-Ortí, A., Arrébola, J. R., Muñoz, B., Chueca, L. J., & Madeira, M. J. (2015). Molecular phylogeny of the western Palaearctic Helicoidea (Gastropoda, Stylommatophora). Molecular Phylogenetics and Evolution, 83, 99–117. 10.1016/j.ympev.2014.11.014

Rivkin, L. R., Santangelo, J. S., Alberti, M., Aronson, M. F. J., de Keyzer, C. W., Diamond, S. E., Fortin, M.-J., Frazee, L. J., Gorton, A. J., Hendry, A. P., Liu, Y., Losos, J. B., MacIvor, J. S., Martin, R. A., McDonnell, M. J., Miles, L. S., Munshi-South, J., Ness, R. W., Newman, A. E. M., … Johnson, M. T. J. (2019). A roadmap for urban evolutionary ecology. Evolutionary Applications, 12(3), 384–398. 10.1111/eva.12734

Rodriguez-Cabal, M. A., Gibbons, T. C., Schulte, P. M., Barrios-Garcia, M. N., & Crutsinger, G. M. (2015). Comparing functional similarity between a native and an alien slug in temperate rain forests of British Columbia. NeoBiota, 25, 1–14. 10.3897/neobiota.25.8316

Simonová, J., Simon, O. P., Kapic, Š., Nehasil, L., & Horsák, M. (2016). Medium-sized forest snails survive passage through birds’ digestive tract and adhere strongly to birds’ legs: More evidence for passive dispersal mechanisms. Journal of Molluscan Studies, 82(3), 422–426. 10.1093/mollus/eyw005

Smith, S. D. A., & Nimbs, M. J. (2022). Citizen Scientists Record Significant Range Extensions for Tropical Sea Slug Species in Subtropical Eastern Australia. Diversity, 14(4), Article 4. 10.3390/d14040244

Soltis, D. E., Gitzendanner, M. A., Strenge, D. D., & Soltis, P. S. (1997). Chloroplast DNA intraspecific phylogeography of plants from the Pacific Northwest of North America. Plant Systematics and Evolution, 206(1–4), 353–373. 10.1007/BF00987957

Tamura, K., & Nei, M. (1993). Estimation of the number of nucleotide substitutions in the control region of mitochondrial DNA in humans and chimpanzees. Molecular Biology and Evolution, 10, 512–526.

Vendetti, J., Lee, C., & LaFollette, P. (2018). Five New Records of Introduced Terrestrial Gastropods in Southern California Discovered by Citizen Science. American Malacological Bulletin, 36, 232. 10.4003/006.036.0204

von Proschwitz, T., Reise, H., Schlitt, B., & Breugelmans, K. (2017). Records of the slugs Ariolimax columbianus (Ariolimacidae) and Prophysaon foliolatum (Arionidae) imported into Sweden. Folia Malacologica, 25(4), 267–271. 10.12657/folmal.025.023

Wade, C. M., Mordan, P. B., & Naggs, F. (2006). Evolutionary relationships among the Pulmonate land snails and slugs (Pulmonata, Stylommatophora): LAND SNAIL PHYLOGENY. Biological Journal of the Linnean Society, 87(4), 593–610. 10.1111/j.1095-8312.2006.00596.x

Wilke, T., & Duncan, N. (2004). Phylogeographical patterns in the American Pacific Northwest: Lessons from the arionid slug Prophysaon coeruleum. Molecular Ecology, 13(8), 2303–2315. 10.1111/j.1365-294X.2004.02234.x

Williams, B.D., Schrank, B., Huynh, C., Shownkeen, R., & Waterston, R.H. (1992). A genetic mapping system in Caenorhabditis elegans based on polymorphic sequence tagged sites. Genetics, 131:15.

Wilson, J. S., Pan, A. D., General, D. E. M., & Koch, J. B. (2020). More eyes on the prize: An observation of a very rare, threatened species of Philippine Bumble bee, Bombus irisanensis, on iNaturalist and the importance of citizen science in conservation biology. Journal of Insect Conservation, 24(4), 727–729. 10.1007/s10841-020-00233-3

